# Towards Understanding Comprehensive Morphometric Changes and Its Correlation with Cognition and Exposure to Fighting in Active Professional Boxers

**DOI:** 10.1101/2021.09.25.461817

**Authors:** Virendra R. Mishra, Xiaowei Zhuang, Karthik R. Sreenivasan, Dietmar Cordes, Aaron Ritter, Sarah J. Banks, Charles Bernick

**Author notes:** Corresponding Author: Virendra Mishra, Ph.D. Associate Staff Cleveland Clinic Lou Ruvo Center for Brain Health 888 W. Bonneville Avenue Las Vegas, NV 89106, USA.

## Abstract

Professional athletes exposed to repetitive head impacts are at increased risk for developing a progressive neurological syndrome known as traumatic encephalopathy syndrome and neuropathology seen on autopsy called chronic traumatic encephalopathy (CTE). The early identification of individuals at increased risk for CTE is important and the search for biomarkers is underway. In this study, we utilized data from a large cohort study to compare differences in regional brain volumes, cortical thickness, voxel-based morphometric (VBM)-derived measures, and graph-theoretical measures derived from large-scale topographical maps in active professional boxers. We compared the above morphometric measures between active professional boxers with low cognitive scores (impaired boxers) and active professional boxers with intact cognitive scores (nonimpaired boxers). The cognitive scores were evaluated through neuropsychological evaluation. As an exploratory analysis, we also examined the power of various machine-learning algorithms to identify impaired and nonimpaired boxers using both group-level regression-driven analysis and previously identified hypothesis-driven cortical thickness and volumetric measures. We found significant group-level differences between impaired and nonimpaired boxers in cortical thickness in a single brain region (right precuneus), differences in VBM-derived gray matter density encompassing the caudate, putamen, and thalamus; and white matter density encompassing the right paracentral lobule, but no differences in any graph-theoretical network properties. Additionally, we found that *a priori* hypothesis-driven T1-derived cortical thickness and volumetric analysis performed better than traditional regression-based analysis. Overall, this study suggests that neuroanatomical differences exist between impaired and nonimpaired active professional boxers, and that hypothesis-driven techniques are likely necessary to become reliable biomarkers.

## 1. INTRODUCTION

Repetitive head impacts (RHI) are the primary risk factor associated with the pathological accumulation of phosphorylated tau in neurons and astroglia known as chronic traumatic encephalopathy (CTE) (McKee et al., 2016). CTE has been observed at autopsy in professional athletes from a variety of different contact sports and military veterans (Maroon et al., 2015). However, CTE cannot be diagnosed currently in a living brain. Katz et al. recently proposed a clinical criterion for Traumatic Encephalopathy Syndrome (TES) (Katz et al., 2021) which is based on retrospective clinicopathological studies of individuals with CTE, and may assist clinicians in the diagnosis of CTE in a living brain.

Progressive memory loss and executive dysfunction such as slower processing speed and difficulty in completing complex attentional tasks are some criteria suggestive of CTE pathology (Katz et al., 2021), and such executive dysfunctions have been reported in neuropsychological studies of professional fighters (Bernick and Banks, 2013; Förstl et al., 2010; Heilbronner et al., 2009). However, epidemiological studies suggest that only 10-20% of professional boxers suffer from significant and persistent neurological deficits (Förstl et al., 2010). Hence identification of participants with a significant risk of developing neuropathological changes will allow fighters to make informed decisions about their ongoing careers.

Significant efforts have been made to understand the neuroanatomical correlates of the cognitive scores evaluated through neuropsychological assessments through Positron Emission Tomography (PET) and Magnetic resonance imaging (MRI) studies. Establishing these correlates could be important because they could be further developed as in-vivo biomarkers. MRI techniques are likely better suited than PET because MRI can inform about both the structural and functional brain organization due to RHI. For instance, T1-weighted MRI studies in participants exposed to RHI such as veterans, fighters, and football players have shown widespread volumetric deficits involving the thalamus, fusiform gyrus, and ventromedial prefrontal cortex (Bernick et al., 2015; Bigler, 2013; Gooijers et al., 2013; Lopez-Larson et al., 2013; Mishra et al., 2017; Montenigro et al., 2015; Ng et al., 2014). Several neuroimaging studies utilizing only T1-weighted structural MRI has shown deficits in gray-matter (GM) volumes of the thalamus, ventromedial prefrontal cortices, fusiform gyrus, hippocampus, medial temporal lobes, and frontal lobes (Bernick et al., 2015; Bernick and Banks, 2013; Bigler, 2013; Ng et al., 2014) in professional fighters. Similarly, diffusion-weighted MRI (dMRI) studies have reported widespread deficits in the organization of major white-matter (WM) fibers such as corpus-callosum and temporo-occipital WM tracts (Hulkower et al., 2013; Mishra et al., 2017; Ng et al., 2014; Shin et al., 2014; Wintermark et al., 2015; Zhang et al., 2003). Long-range functional connectivity deficits supplementing WM connectivity loss in fighters (Mishra et al., 2019) have also been observed recently through resting-state functional MRI (rsfMRI) study (Zhuang et al., 2020). Likewise, PET studies have indicated perfusion deficits with lower cerebral blood flow in the thalamus, cingulate gyri, temporal lobes, and cuneus (Eierud et al., 2014; Inga K Koerte et al., 2016; Mishra et al., 2018). Furthermore, these deficits in structural, perfusion, fMRI, or dMRI measures were shown to be correlated with exposure to fighting and neurocognitive assessments (Bernick et al., 2015; Ge et al., 2009; Inga K Koerte et al., 2016; Liu et al., 2013; Mishra et al., 2018, 2017; Orrison et al., 2009; Zhuang et al., 2020).

In addition to volumetric measures, T1-weighted MRI can also be used to derive cortical thickness (Fischl and Dale, 2000), compare them (Lin et al., 2017; Pettigrew et al., 2016; Querbes et al., 2009; Tessitore et al., 2016), and investigate if cortical thickness measures are associated with exposure to fighting or cognitive scores observed clinically. Additionally, coordinated changes in cortical thickness could also be evaluated through graph-theoretical measures that may inform about early neurodegeneration. Graph-theoretical measures provide quantitative insight into network parameters which have a strong influence on the dynamics of the network with pathology (Bassett and Bullmore, 2006). For instance, coordinated variation of regional cortical thickness has been reported in neurodegenerative disorders such as Alzheimer’s disease (AD) (He et al., 2008; Xie and He, 2011), Parkinson’s disease (PD) (Yadav et al., 2016), military veterans (Proessl et al., 2020), and depression (Wang et al., 2016). Recent studies have shown deficits in graph-theoretical informed properties during both neurodevelopment (Ball et al., 2014; Huang et al., 2015; Koenis et al., 2017; van den Heuvel et al., 2015) and neurodegeneration (delEtoile and Adeli, 2017; Luo et al., 2015; Mears and Pollard, 2016; Mishra et al., 2020; Utianski et al., 2016). However, it is currently unknown whether there is a coordinated change in cortical thickness in active professional boxers that is correlated with cognitive scores evaluated through neuropsychological evaluations and exposure to fighting. Understanding of these coordinated changes, if any, might help determine if there is preferential topological damage due to RHI.

Hence, in this study, we used the data from the Professional Athletes Brain Health Study (PABHS) (Bernick et al., 2013), and compared cortical thickness and volumetric measures between active professional boxers with low cognitive scores (impaired boxers) and active professional boxers with intact cognitive scores (nonimpaired boxers). We used conventional group-wise cortical thickness statistical comparison techniques and voxel-based-morphometry (VBM) techniques to perform such regression-driven group-level investigations. We also investigated the correlation between T1-weighted MRI-derived measures of cortical thickness and volumetric measures with exposure to fighting and neuropsychological assessments. Furthermore, we also used graph-theoretical measures to identify whether there are large-scale topological differences in the same cohort of active professional boxers.

As an exploratory analysis, we also tested the individual predictive ability of these group-level regression-driven differences with different machine-learning (ML) algorithms along with the predictive ability of previously identified (Mishra et al., 2017) hypothesis-driven cortical thickness and volumetric measures to understand if these T1-derived measures can identify impaired and nonimpaired boxers. The motivation of this exploratory analysis was to identify the most robust T1-derived measure that can serve as biomarkers to identify impaired active professional boxer early in their career. Of note, we only extracted T1-derived measures from the previous study (Mishra et al., 2017) as standardization of both dMRI and rsfMRI measures is still under active investigation.

## 2. MATERIALS AND METHODS

PABHS (Bernick et al., 2013) is a study of active professional fighters (boxers, MMA, and bull riders) that has been following professional athletes for more than 10 years. PABHS was approved by the institutional review board of Cleveland Clinic and all the participants provided informed written consent. The protocol was explained to all the participants and was performed according to the Declaration of Helsinki guidelines and the Belmont Report.

### 2.1. Recruitment criteria of active professional fighters

464 active professional fighters were recruited at our center between 2011-2015. All participants were licensed for professional boxing or mixed martial arts (MMA), fluent in English, and a good read at least at a 4^th^-grade level. Fighters with a sanctioned competition within 45 days of the visit were excluded. Information about educational attainment and detailed information on prior involvement in other contact sports and professional fighting were recorded for most of the participants.

### 2.2. MRI Data Acquisition

3T Verio Siemens MRI scanner with a 32-channel head coil was used to acquire 3D magnetization prepared rapid gradient echo (MPRAGE) T1-weighted scan with the following imaging parameters: field of view (FOV) =256×256×160; voxel size=1×1×1.2mm^3^; TR=2300ms, TE=2.98ms; TI=900ms and flip angle=9°. The total imaging scan time was 9 minutes.

### 2.3. Cortical surface reconstruction using FreeSurfer

The cortical surface was reconstructed for each participant using the FreeSurfer 6.0 pipeline (Fischl et al., 2004; Fischl and Dale, 2000). Briefly, each T1-weighted image was spatially, and intensity normalized to Talairach atlas. Volumetric segmentation and subcortical labeling were then performed on the normalized images. The GM and WM boundaries were then automatically identified and were reconstructed into a mesh of over 150000 tessellated vertices for surface measures at each point that was consistent across all participants. Gyral anatomy was then aligned to a standard spherical template using surface convexity and curvature measures.

### 2.4. Quality control analysis of cortical reconstruction data

An estimate of contrast-to-noise ratio (CNR) in the WM was computed for every participant using the FreeSurfer’s QA tools (https://surfer.nmr.mgh.harvard.edu/fswiki/QATools). Only data with a CNR >16 (https://surfer.nmr.mgh.harvard.edu/pub/dist/freesurfer/tutorial_packages/OSX/freesurfer/bin/wm-anat-snr) was utilized for further analysis. We performed a visual confirmation of choice of this CNR by verification of cortical reconstructions in participants following the procedure outlined in Fjell et al. (Fjell et al., 2009).

The above-mentioned procedure yielded 367 fighters that had acceptable cortical reconstructed data.

### 2.5. Classification of fighters into impaired and nonimpaired fighters

Similar to our previous studies (Mishra et al., 2017; Zhuang et al., 2020), we categorized our fighters into impaired and nonimpaired groups using the neuropsychological test scores of processing speed and psychomotor speed. Importantly, we did not explore whether the cognitive scores evaluated through these neuropsychological evaluations caused difficulty in the day-to-day functioning of the enrolled fighters. Cognitive tests were completed using a computer in a quiet room, supervised by a researcher. CNS Vital Signs was used to administer standardized cognitive tests (Gualtieri and Johnson, 2006). We used two tests from their battery: Finger Tapping and Digit Symbol Coding. From these tests, we obtained processing speed (total correct on a Digit Symbol Coding task) and psychomotor speed (combining Digit Symbol result and average Finger Tapping on each hand) for all the participants. The clinical scores hence obtained on all participants were standardized by converting to z-scores. The z-scores of all participants were obtained by comparing the values to an age-matched healthy controls normative group that is proprietary to CNS. Three fighters were unable to finish the required neuropsychological assessments.

Fighters that had either or both standardized processing speed and standardized psychomotor speed two standard deviations below the mean were classified as impaired (Schinka et al., 2010). The remaining fighters formed the nonimpaired group yielding a total of 116 impaired fighters and 248 nonimpaired fighters. Since there is a significant effect of sex on brain volumes in both healthy aging (Wang et al., 2019), and RHI (Bennett et al., 2020), we only utilized male professional fighters in our cohort. Furthermore, since boxers and MMA have distinct relationships between exposure to fighting and neuroanatomic findings (Shin et al., 2014), we only included male professional boxers in our cohort to reduce the bias due to the type of professional fighting. This constraint on sex and fighting style yielded 79 impaired and 136 nonimpaired male boxers. Numbers and years of professional fighting were recorded only on 72 impaired and 130 nonimpaired boxers in this cohort. Since exploring ML techniques was one of the goals of this study, and the ML algorithms are biased with unequal sample sizes (Max and Johnson, 2013), we selected 72 nonimpaired boxers at random that were matched for independent variables such as years of education, age, and number and years of professional fighting.

The demographics and neuropsychological scores for the final set of 72 impaired and 72 nonimpaired active professional male boxers used in this study are tabulated in Table 1.

**Table 1:**
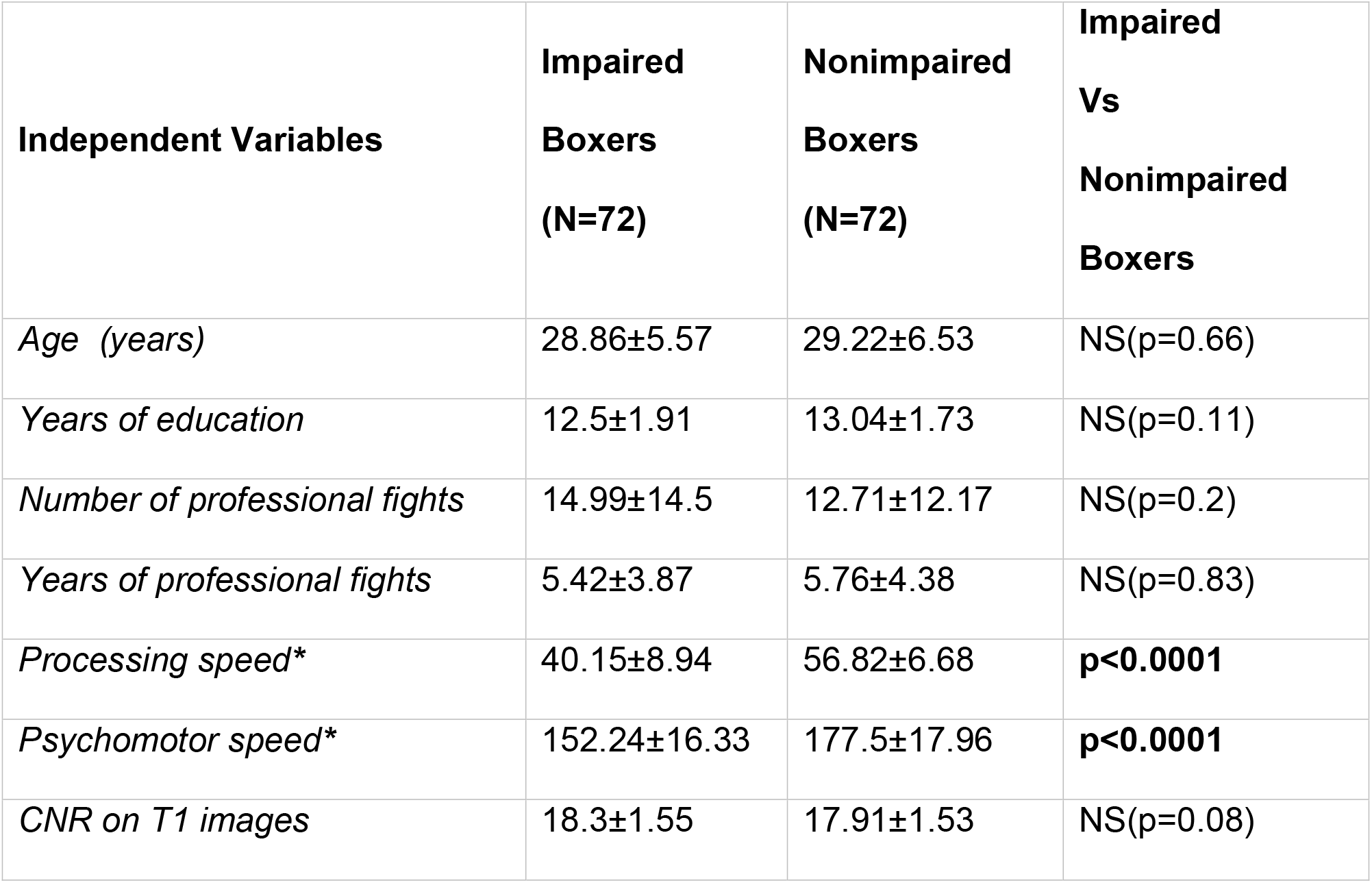
Various demographics along with the neuropsychological assessment test results of all the participants are shown along with their mean±SD. Results of pairwise statistical comparisons are also shown as p-values, represented by the letter “p”. NS: Non-significant; NA: Not-applicable.

### 2.6. Cortical thickness analysis

The final thickness map of each participant from FreeSurfer was coregistered to a standard template (fsaverage) and was used for cortical thickness analysis. Cortical thickness and volume measures were extracted for regions specified in the Desikan-Killiany atlas (Desikan et al., 2006) in FreeSurfer.

### 2.7. VBM analysis

VBM analysis was performed using Diffeomorphic Anatomical Registration Through Exponential Lie Algebra (DARTEL) toolbox (Ashburner, 2007) to compare modulated gray matter density (GMD) and white matter density (WMD) between impaired and nonimpaired boxers. Briefly, the steps can be summarized as follows: (1) T1-weighted images were segmented using standard unified segmentation model in SPM12 to produce GM, WM, and cerebrospinal fluid (CSF) probability maps in MNI152 space; (2) the study-specific GM and WM templates were created from the segmented GM and WM images by selecting all impaired and nonimpaired boxers to stay unbiased to population distribution; (3) an initial affine registration of both GM and WM templates to tissue probability maps in MNI152 space was then performed; (4) both affine registered GM and WM templates were nonlinearly warped to GM and WM templates in MNI152 space respectively, and were used in the modulation step in order to ensure that the relative volumes of GM and WM were preserved following the spatial normalization (1mm^3^ isotropic resolution) by the Jacobian determinant of the deformation field; and (5) spatial smoothing was finally performed on the modulated and normalized GM and WM images with a 10mm full width at half maximum (FWHM) isotropic Gaussian kernel and were used to statistically compare modulated GMD and WMD between the two groups. Of note, WMD obtained using DARTEL encompasses both the WM and the subcortical brain regions.

### 2.8. Graph-theoretical analysis

The large-scale topological structure of a network can be estimated with various graph-theoretical measures. Since we wanted to investigate topological disruptions between impaired and nonimpaired boxers at a group level, we generated a group level symmetric connectivity matrix (He et al., 2008). The nodes and the edges of the connectivity matrix in this study were defined as follows:

#### Node definition

Each anatomical region in the Desikan-Killiany atlas (Desikan et al., 2006) was defined as a node of the cortical network yielding 34 nodes in each hemisphere for each participant. Table 2 and the top panel of Fig.3 shows the nodes used for this study.

**Table 2:** The regions from FreeSurfer used in the current study are tabulated. Reg ID, R, and L represent the region identification number, right hemisphere, and left hemisphere respectively.

| Node name in the atlas | Reg ID # | Node name | Reg ID # | Node name |
| --- | --- | --- | --- | --- |
| bankssts | 1 | *'R.Bankssts'* | 35 | *'L.Bankssts'* |
| caudal anterior cingulate | 2 | *'R.CaudAntCing'* | 36 | *'L.CaudAntCing'* |
| caudal middle frontal | 3 | *'R.CaudMidFron'* | 37 | *'L.CaudMidFron'* |
| cuneus | 4 | *'R.Cun'* | 38 | *'L.Cun'* |
| entorhinal | 5 | *'R.Ent'* | 39 | *'L.Ent'* |
| fusiform | 6 | *'R.Fus'* | 40 | *'L.Fus'* |
| inferior parietal | 7 | *'R.InfPar'* | 41 | *'L.InfPar'* |
| inferior temporal | 8 | *'R.InfTemp'* | 42 | *'L.InfTemp'* |
| isthmus cingulate | 9 | *'R.IsthCing'* | 43 | *'L.IsthCing'* |
| lateral occipital | 10 | *'R.LatOcc'* | 44 | *'L.LatOcc'* |
| lateral orbito frontal | 11 | *'R.LatOrbFron'* | 45 | *'L.LatOrbFron'* |
| lingual | 12 | *'R.Ling'* | 46 | *'L.Ling'* |
| medial orbito frontal | 13 | *'R.MedOrbFron'* | 47 | *'L.MedOrbFron'* |
| middle temporal | 14 | *'R.MidTemp'* | 48 | *'L.MidTemp'* |
| parahippocampal | 15 | *'R.ParaHipp'* | 49 | *'L.ParaHipp'* |
| paracentral | 16 | *'R.ParaCent'* | 50 | *'L.ParaCent'* |
| pars opercularis | 17 | *'R.ParsOper'* | 51 | *'L.ParsOper'* |
| pars orbitalis | 18 | *'R.ParsOrb'* | 52 | *'L.ParsOrb'* |
| pars triangularis | 19 | *'R.ParsTri'* | 53 | *'L.ParsTri'* |
| pericalcarine | 20 | *'R.PeriCal'* | 54 | *'L.PeriCal'* |
| postcentral | 21 | *'R.PostCen'* | 55 | *'L.PostCen'* |
| posterior cingulate | 22 | *'R.PostCing'* | 56 | *'L.PostCing'* |
| precentral | 23 | *'R.PreCen'* | 57 | *'L.PreCen'* |
| precuneus | 24 | *'R.PreCun'* | 58 | *'L.PreCun'* |
| rostral anterior cingulate | 25 | *'R.RostAntCing'* | 59 | *'L.RostAntCing'* |
| rostral middle frontal | 26 | *'R.RostMidFron'* | 60 | *'L.RostMidFron'* |
| superior frontal | 27 | *'R.SupFron'* | 61 | *'L.SupFron'* |
| superior parietal | 28 | *'R.SupPar'* | 62 | *'L.SupPar'* |
| superior temporal | 29 | *'R.SupTemp'* | 63 | *'L.SupTemp'* |
| supramarginal | 30 | *'R.Supramarg'* | 64 | *'L.Supramarg'* |
| frontal pole | 31 | *'R.FronPole'* | 65 | *'L.FronPole'* |
| temporal pole | 32 | *'R.TempPole'* | 66 | *'L.TempPole'* |
| transverse temporal | 33 | *'R.TransTemp'* | 67 | *'L.TransTemp'* |
| insula | 34 | *'R.Ins'* | 68 | *'L.Ins'* |

#### Edge definition

The edges between the above-defined nodes in this study were represented by the absolute partial correlation of the cortical thickness between each node (He et al., 2008), as partial correlation represents the correlation between any two nodes after regressing the effect of other nodes. The procedure to generate group-level connectivity matrices is similar to the procedure by He et al. (He et al., 2008). Briefly, the steps can be summarized as follows. First, using the tools in FreeSurfer, we extracted the average cortical thickness of each node for each participant. The cortical thickness within each group was then regressed for age, years of education, average left cortical thickness of each participant, average right cortical thickness of each participant, and total intracranial volume; and the residuals were used as a surrogate for cortical thickness for each participant within each group (He et al., 2008; Phillips et al., 2015). Then, we generated the connectivity matrices by computing the interregional correlation C(*i,j*) (*i,j = 1 to 68 nodes)* between each node. This procedure yielded a 68×68 symmetric connectivity matrix for the impaired and nonimpaired boxers group. Since the study focused on monotonic coordinated change in cortical thickness between the groups, an absolute partial Spearman’s rank correlation (ρ) was then computed between the regressed cortical thicknesses of each node within the group. To control for the effect of possible different absolute correlations between the two groups, a sparsity threshold was applied that guarantees the number of edges or the wiring cost in the graph of two groups to be equal (Achard and Bullmore, 2007; Stam et al., 2007; Telesford et al., 2010). This procedure allows for a fair comparison of alterations of topological organization between the groups. In the absence of any known single sparsity threshold, a range of sparsity threshold (He et al., 2008) from 5% to 40% in steps of 1% was applied to generate 36 binarized connectivity matrices for the two groups.

We computed various graph-theoretical measures of each group at each sparsity threshold either using custom Matlab® scripts, GRETNA (J. Wang et al., 2015), or Brain Connectivity Toolbox (https://sites.google.com/site/bctnet/). Various global measures such as normalized clustering coefficient (γ), normalized path length (λ), small-worldness (σ), local efficiency, and global efficiency (Latora and Marchiori, 2003; Rubinov and Sporns, 2010; Watts and Strogatz, 1998) were computed at each sparsity threshold. We also computed the nodal degree (Wang et al., 2010) of each node for the fully connected graph.

Briefly, the measures used in this study could be summarized as follows:

#### Nodal Degree

Degree of a node is the simplest graph-theoretical metric that describes the connectivity of a node with the rest of the nodes in the network. The degree of each node *i* was computed as:

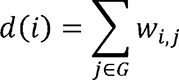

where *d(i)* represents the degree of node *i, G* represents the graph, *j* represents the other nodes of the graph *G*, and *w_i,j_* represents the edge weight between the nodes *i* and *j* in the connectivity matrix. The edge weights (*w_i,j_)* of the network were binarized in our study and varied depending upon the sparsity threshold used.

#### Global Efficiency

The global efficiency (*E_glob_ (G)*) of the graph *G* measures the efficiency of the parallel information transfer in the network. *E_glob_(G)* was calculated as:

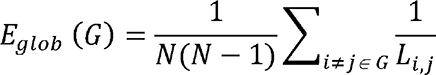

where *N* represents the total number of nodes in the graph and *L_i,j_* is the shortest path length between the nodes *i* and *j* which was computed as the inverse of the edge weight (*w_i,j_*).

#### Local Efficiency

The local efficiency (*E_loc_(G))* of the graph *G* represents the efficiency of the information flow among first neighbors of the node *i* when it is removed. *E_loc_(G)* provides information about network resilience to fault tolerant. *E_loc_(G)* was calculated as:

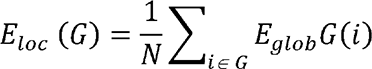

where *G_i_* is the subgraph of the nearest neighbors of node *i*.

#### Normalized clustering coefficient

The extent of local interconnectivity or network segregation of each network graph (*G)* was computed as network clustering coefficient *(Cp)* computed as:

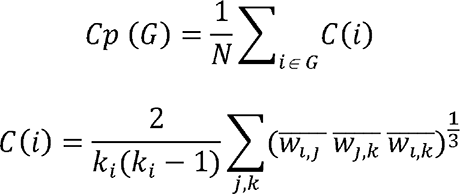

where *C(i)* is the clustering coefficient of node *i*, *k_i_* is the degree of node *i*, and 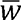 is the weight scaled by the mean of all weights. As can be observed, *C(i)=*0 if the nodes are isolated or *k_i_=*0 or 1. Normalized clustering coefficient (γ) was computed by normalizing each graph’s 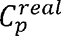 *(G)* by 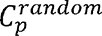 *(G)* where 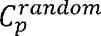*(G)* is the mean clustering coefficient of 100 degree-matched random networks, and 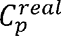 *(G)* is the clustering coefficient of the real graph *G*.

#### Normalized path length

Network integration of a graph *G* measures the network’s ability to transfer information in parallel. It was computed as network shortest path length *(Lp)* as:

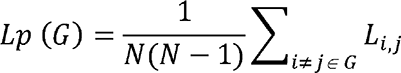

where *L_i,j_* is defined as the length of the path for nodes *i* and *j* with the shortest length, and length of each edge was defined as the inverse of the edge weight *(w_i,j_)*. Normalized shortest path length (λ) was computed by normalizing each graph’s 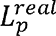 *(G)* by 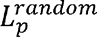 *(G)*.

#### Small-worldness

Small-worldness (σ) of the graph *G* is defined as a ratio between normalized clustering coefficient and normalized shortest path length as σ=γ/λ. For a network to be small-world, γ>>1 and λ≍1 (He et al., 2008; Watts and Strogatz, 1998). It represents both efficient network segregation and network integration (Rubinov and Sporns, 2010).

### 2.9. Statistical analysis

#### Vertex-wise group analysis to test for group differences and correlation with exposure to fighting and cognitive scores

A nonparametric approach using FSL’s permutation analysis of the linear model (PALM) (Winkler et al., 2016) with threshold-free cluster enhancement (TFCE) (Smith and Nichols, 2009) was used to perform a vertex-wise comparison of the cortical thickness between the groups. Age, years of education, average left cortical thickness of each participant, average right cortical thickness of each participant, and total intracranial volume were used as covariates of no interest. Vertex-wise correlation of both exposures to fighting and cognitive measures with the cortical thicknesses within each group was also conducted using PALM after controlling for the nuisance covariates. Significance for both vertex-wise group difference and correlation between exposure to fighting and cognitive scores with vertex-wise cortical thickness was established at p_corr_<0.05.

#### VBM analysis to test for group differences and correlation with exposure to fighting and cognitive scores

The nonparametric approach using FSL’s PALM with TFCE with the same design and contrast matrix as described above was also used to perform VBM analysis to compare differences in GMD and WMD between the groups. VBM analysis to test the correlation between GMD and WMD with both exposures to fighting and cognitive scores within each group was also conducted using PALM after controlling for the nuisance covariates. Significance for both voxel-wise group difference and correlation between exposure to fighting and cognitive scores with voxel-wise GMD and WMD was established at p_corr_<0.05.

#### Statistical analysis of graph-theoretical derived metrics to test for group differences

A nonparametric permutation test was used to determine the statistical significance of the differences in both global and local graph-theoretical derived metrics between the two groups. To test the null hypothesis that the observed group differences could occur by chance, a null model was generated as outlined in He et al. (He et al., 2008). First, each participant’s set of regional cortical thickness measures were randomly reallocated to either group and a partial correlation matrix was recomputed for each randomized group. The matrix thus obtained was then binarized at the same sparsity threshold and the same network parameters were calculated as the real networks. Finally, a group difference of the network parameters was computed. This procedure of random permutation was repeated 1000 times and the 95^th^ percentile of the distribution was used as the cutoff for the null hypothesis with a probability of type I error of 0.05. The above steps were repeated at each sparsity threshold value for the brain networks. Significance for both group differences of various graph-theoretical measures and correlation between exposure to fighting and cognitive scores with various graph-theoretical measures was established at p_corr_<0.05.

Of note, each technique (vertex-wise, VBM, and graph-theoretical measures) were treated independently of each other, and no multiple comparison corrections were performed between the three techniques.

### 2.10. Predicting impaired and nonimpaired active professional boxers

We utilized four widely-used ML algorithms, namely radial basis functional networks (RBFN) (Broomhead and Lowe, 1988), support vector machine (SVM) with linear and nonlinear (RBF) kernel (Ben-Hur et al., 2008), and random forest (Liaw and Wiener, 2002) to test the exploratory goal of identifying impaired and nonimpaired active professional boxers.

Following feature sets were used for every ML technique: (a) all FreeSurfer-derived region-of-interest based cortical thickness and volume measures, (b) regional cortical thickness with significant (p_corr_<0.05) univariate differences between impaired and nonimpaired boxers, (c) regional GMD and WMD with significant (p_corr_<0.05) univariate differences between impaired and nonimpaired boxers, (d) graph-theoretical measures with significant (p_corr_<0.05) univariate differences between impaired and nonimpaired boxers, (e) regression-derived brain regions significantly correlated with neuropsychological scores or exposure to fighting in univariate vertex-wise analysis, (f) regression-derived GMD and WMD significantly correlated with neuropsychological scores or exposure to fighting in univariate VBM analysis; (g) five *a priori* T1-derived measures namely, left cerebellum white matter volume, right hemisphere cortical volume, left thalamus volume, right pallidum volume, and cortical thickness of right rostral-anterior cingulate(Mishra et al., 2017); and (h) the following combinations of the features described above (*b* and *f*; b and *g*; *f* and *g*; and *b, f* and *g*). The location of the *a priori* T1-derived regions is shown in Fig.6.

80% of data (58 boxers in each group) was used as training dataset and 20% of the dataset (14 boxers in each group) was used as an independent testing dataset. Of note, data from 38/72 impaired boxers and 32/72 nonimpaired boxers utilized in this study were present in the previous study(Mishra et al., 2017) of which 27/58 impaired boxers and 31/58 nonimpaired boxers were in the training dataset, and 7/14 impaired boxers and 5/14 nonimpaired boxers were in the testing dataset. Four benchmark metrics, namely, sensitivity, specificity, area under the receiver operating curve (AUROC), and prediction accuracy were used to identify the optimum ML algorithm with the associated optimized feature parameters that provide the best identification of impaired and nonimpaired boxers. All ML algorithms were optimized for classification in the training dataset using R v1.0 or MATLAB. *caret* package (https://topepo.github.io/caret/) was utilized for the optimization of random forest and SVM parameters. Since the number of cortical thickness and volume measures from all the regions in Desikan-Killiany atlas (feature set, *a*, above) was greater (total features=171; 68 thickness measure, 103 volume measures) than the number of participants in the training dataset (n=116), the features from FreeSurfer were first reduced using Least Absolute Shrinkage and Selection Operator (LASSO) (Tibshirani, 1994). Optimization parameters that maximized the prediction accuracy and AUROC with 10-fold cross-validation in the training dataset were utilized for prediction in the testing dataset.

#### Significance testing of ML approach

Each benchmark measure for every ML algorithm with its associated optimized feature parameters were tested for significance independently of one other in the testing dataset using the nonparametric approach as follows: Each boxer in the testing dataset was randomly assigned as impaired or nonimpaired and the benchmark metric for each ML algorithm with its associated optimized feature parameters obtained from the training dataset, and this procedure was repeated 5000 times. If the 95^th^ percentile of the false-positive error rate of the benchmark metric with random assignments was less than the benchmark metric with true group assignment, then the ML algorithm with its associated optimized feature parameters was considered significant (p_corr_<0.05). Of note, each ML algorithm was treated independently and no statistical comparisons between various ML algorithms were performed.

### 2.11. Data Availability

The de-identified raw MPRAGE data from all the participants along with the scripts used and the participants used for training and testing used for predictive modeling is available to download from https://github.com/diffusionguy/ML_Cortical_Thickness to qualified investigators. To ensure data integrity and appropriate use of the data, a collaborative partnership with the authors is strongly encouraged.

## 3. RESULTS

### 3.1. Statistical results of the demographics and clinical scores

Table 1 outlines the descriptive statistics as mean ± standard deviation (SD) and p-values for the demographics and the cognitive scores among the groups. None of the demographics (age, years of education, number, and years of professional fights) was found to be significantly different between the impaired and nonimpaired groups. Impaired boxers had significantly lower processing speed (p<0.0001) and psychomotor speed (p<0.0001) when compared to nonimpaired boxers, as would be expected since the group was defined based on these criteria. The mean CNR reported by FreeSurfer tools was 18.34±1.55 and 17.91±1.53 in the impaired and nonimpaired boxers group respectively and was not significantly different (p=0.08).

### 2.2. Vertex-wise analysis of cortical thickness

Significantly (p_corr_<0.05) lower cortical thickness (Fig.1) was observed in the right cuneus of impaired boxers (2.06±0.17mm) as compared to nonimpaired boxers (2.09±0.17mm). Cohen’s effect size (d) of this difference was small (0.24±0.003). No brain region was shown to correlate with cognitive scores or years of fighting.

**Fig. 1:**
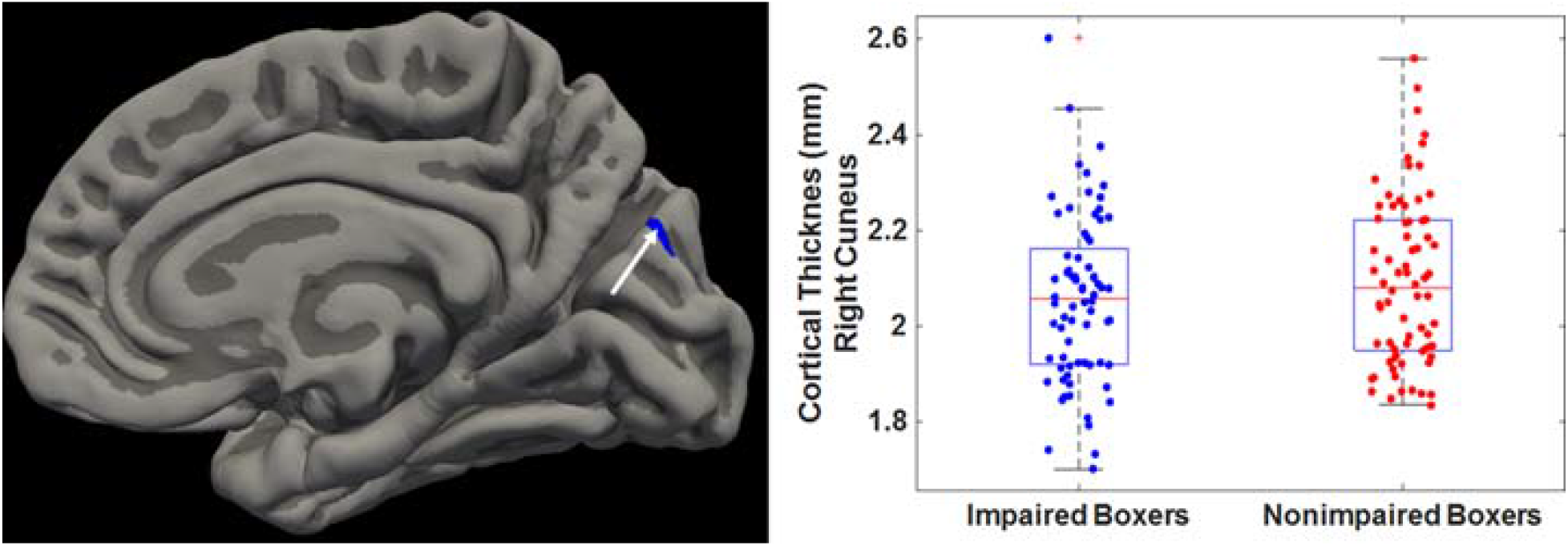
*<u>Left:</u>* Cluster (blue) showing significantly lower cortical thickness in impaired boxers is identified by the arrow and overlaid on the MNI surface. *<u>Right:</u>* Average cortical thickness in the significant cluster is extracted and shown as a boxplot for both groups. Each dot represents average values in the significant cluster for each boxer. Impaired and nonimpaired boxers are shown as blue and red dots respectively.

### 2.3. VBM analysis of GMD and WMD

No significant difference in either whole brain GMD or WMD was observed between the groups. However significantly (p_corr_=0.027±0.03) negative correlation between years of fighting and GMD cluster predominantly encompassing bilateral thalamus, bilateral putamen, and bilateral caudate was observed in the cohort of impaired boxers (r=-0.28±0.01). Impaired boxers also showed a significantly (p_corr_=0.048±0.001) positive correlation between psychomotor speed and WMD cluster predominantly encompassing the right paracentral lobule (r=0.31±0.1). Significantly (p_corr_=0.048±0.002) positive correlation between psychomotor speed and WMD cluster predominantly encompassing left superior frontal gyrus and left precuneus was observed in the cohort of nonimpaired boxers (r=-0.31±0.01). The location of the cluster and the scatterplots of these correlations are shown in Fig.2.

**Fig. 2:**
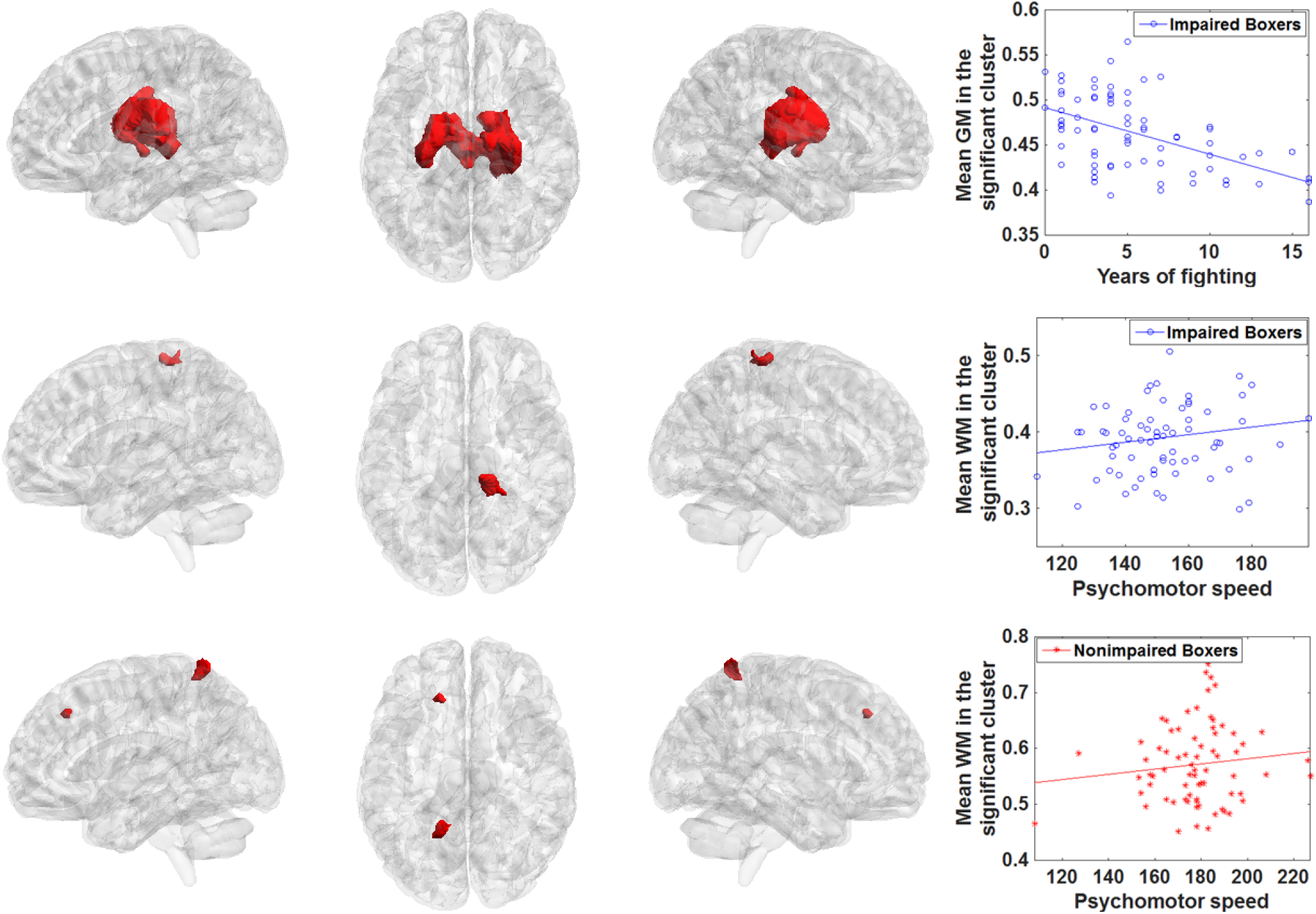
*<u>Top:</u>* Cluster (red) showing a significant correlation between modulated gray-matter (GM) density and years of fighting in impaired boxers. Mean GM density was extracted from each impaired boxer (each blue dot) and plotted as a scatterplot. *<u>Middle:</u>* Cluster (red) showing a significant correlation between modulated white-matter (WM) density and psychomotor speed in impaired boxers. Mean WM density was extracted from each impaired boxer (each blue dot) and plotted as a scatterplot. *<u>Bottom:</u>* Cluster (red) showing a significant correlation between modulated WM density and psychomotor speed in nonimpaired boxers. Mean WM density was extracted from each nonimpaired boxer (each red dot) and plotted as a scatterplot.

### 2.4. Volumetric analysis of T1-derived measures

No significant difference in regional volume measures derived from FreeSurfer was observed between the groups. However significantly (p_corr_=0.045) positive correlation between number of professional fights and volume of the left fusiform gyrus was observed in the cohort of nonimpaired boxers (r=0.3). The location of the left fusiform gyrus and the scatterplot of the correlation is shown in Fig.3.

**Fig. 3:**
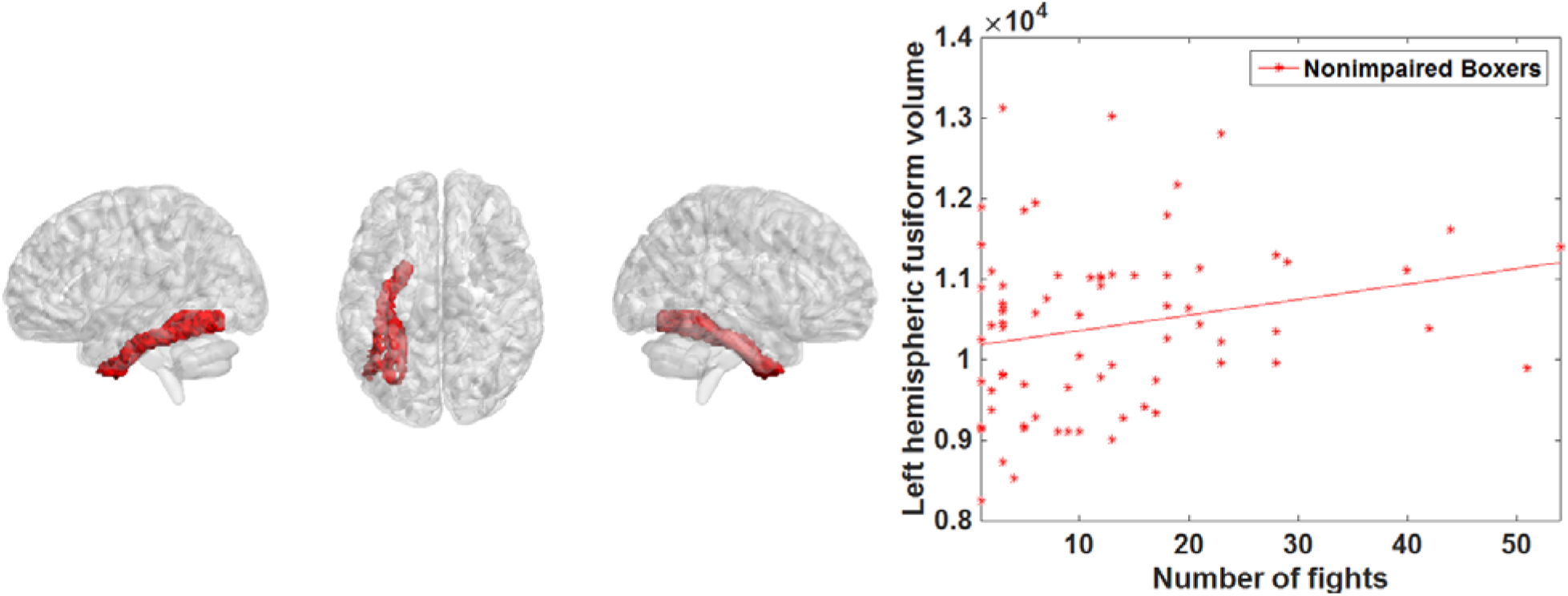
Cluster (red) showing the FreeSurfer parcellated left fusiform gyrus that showed a significant correlation with numbers of fights in nonimpaired boxers. Volume was extracted from each nonimpaired boxer (each red dot) and plotted as a scatterplot.

### 2.5. Global graph-theoretical network results

Both absolute and weighted correlation matrices for both groups are shown in the middle and bottom panel of Fig.4. There was no statistical difference (p_corr_>0.05) in small-worldness, normalized clustering coefficient, and normalized path length between impaired and nonimpaired boxers. We also did not observe any statistical difference (p_corr_>0.05) in global efficiency, local efficiency, and nodal degree between the groups (Fig.5). Furthermore, no correlation between any graph-theoretical derived metrics and cognitive score or exposure to fighting were observed in our cohort.

**Fig. 4:**
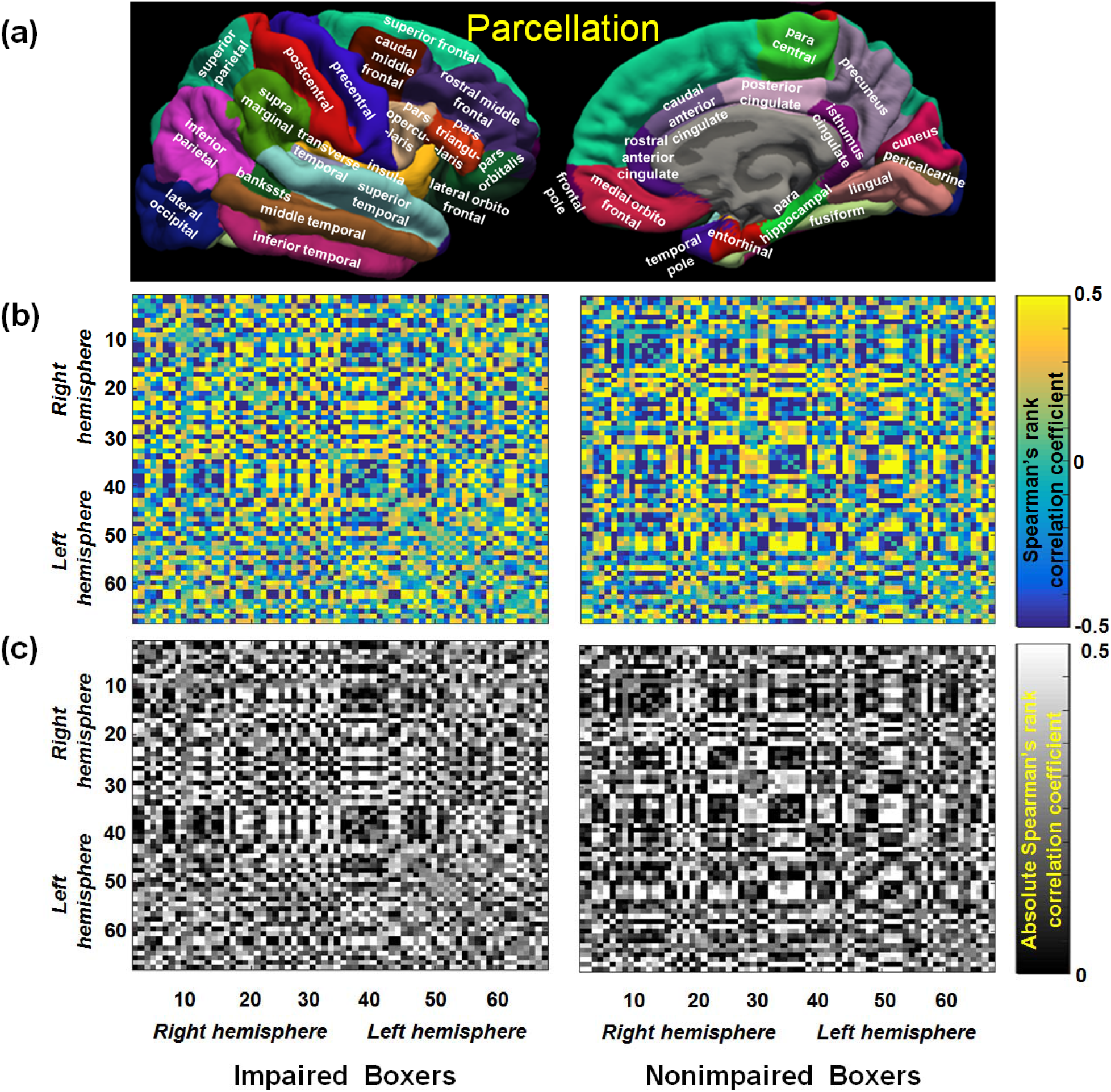
**(a)** The spatial locations of the FreeSurfer regions are shown. **(b)** The Spearman’s rank correlation of the cortical thickness between the FreeSurfer regions for both impaired and nonimpaired fighters group is shown. The color in each entry of the connectivity matrix represents the correlation of the cortical thickness between the respective regions. The color bar represents the range of Spearman’s rank correlation. **(c)** The modulus of the connectivity matrices obtained for both cognitively impaired and nonimpaired active professional fighters in (b) are shown. All the graph-theoretical analysis is done on the binarized matrix of (c). The color bar represents the range of Spearman’s rank correlation. As can be seen, there are visual correlation differences between the groups shown by the orange arrows.

**Fig. 5:**
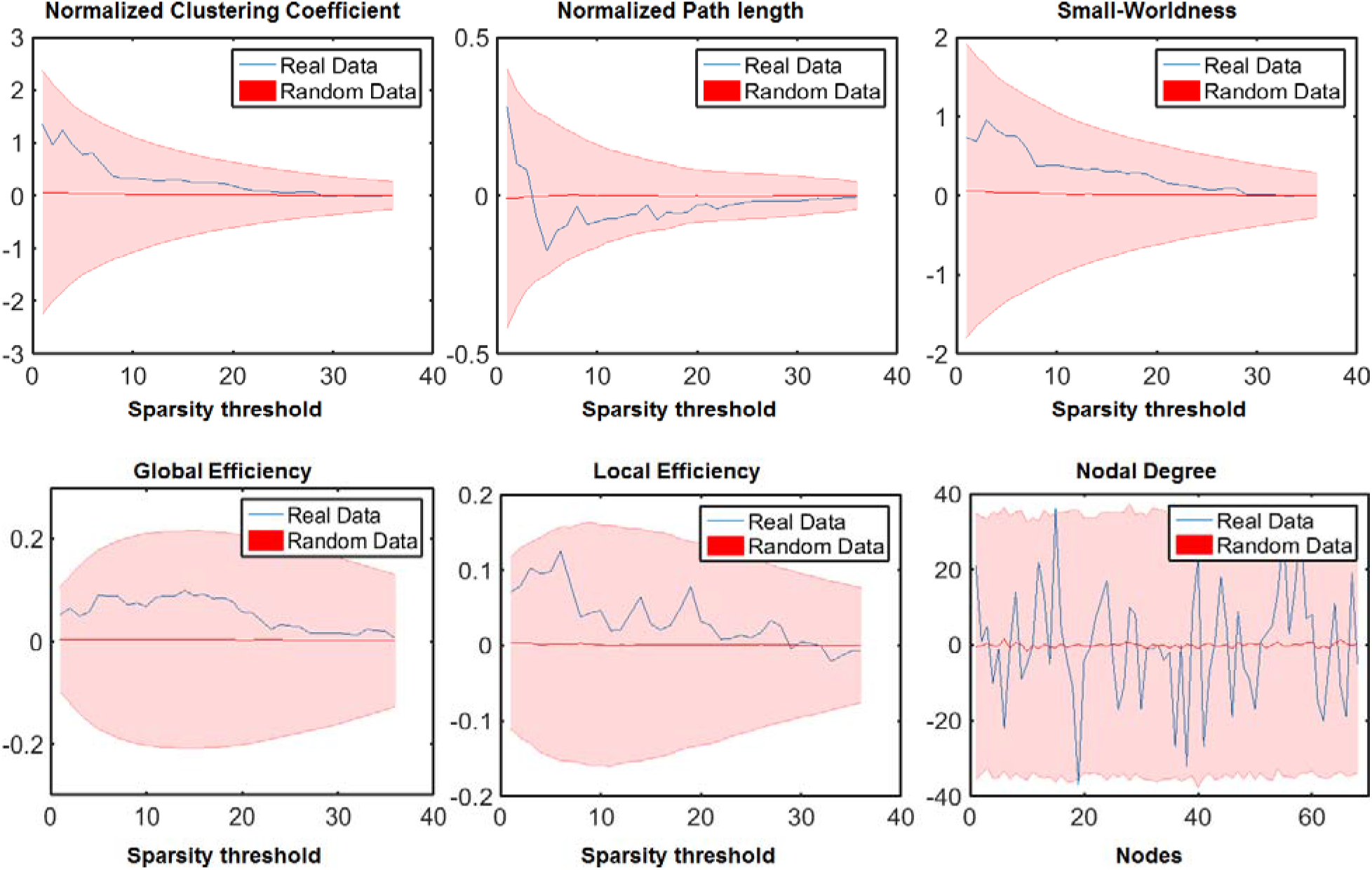
Differences in various graph-theoretical measures against different sparsity threshold (0.05-0.4 in steps of 0.01) for real data (blue) and an average difference of 1000 randomly permuted data (red line) along with the standard deviation of the average difference of 1000 randomly permuted data (pink curve) is shown.

**Fig. 6:**
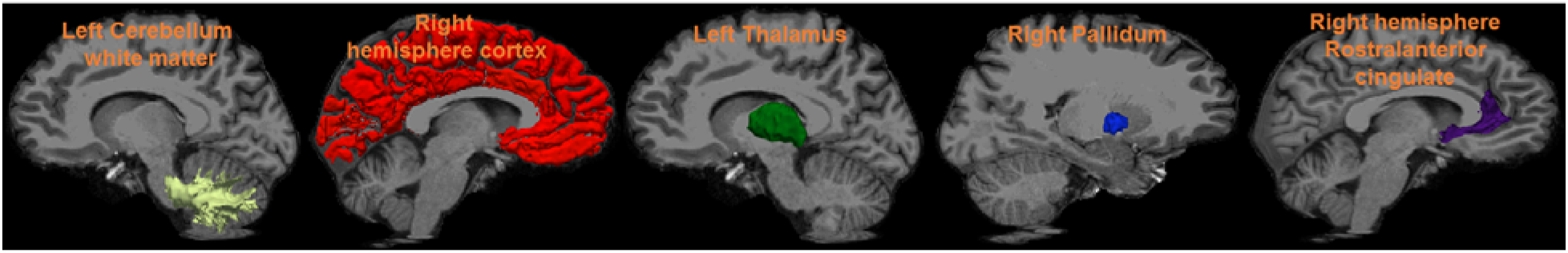
Prior identified brain regions are overlaid on the MNI brain.

### 2.6. Identifying impaired and nonimpaired active professional boxers with features derived from regression-driven methods, hypothesis-driven methods, and combination of regression and hypothesis-driven methods

The benchmark measures with the various feature metrics and ML algorithms are tabulated in Table 3 and are shown graphically in Fig.7.

**Fig. 7:**
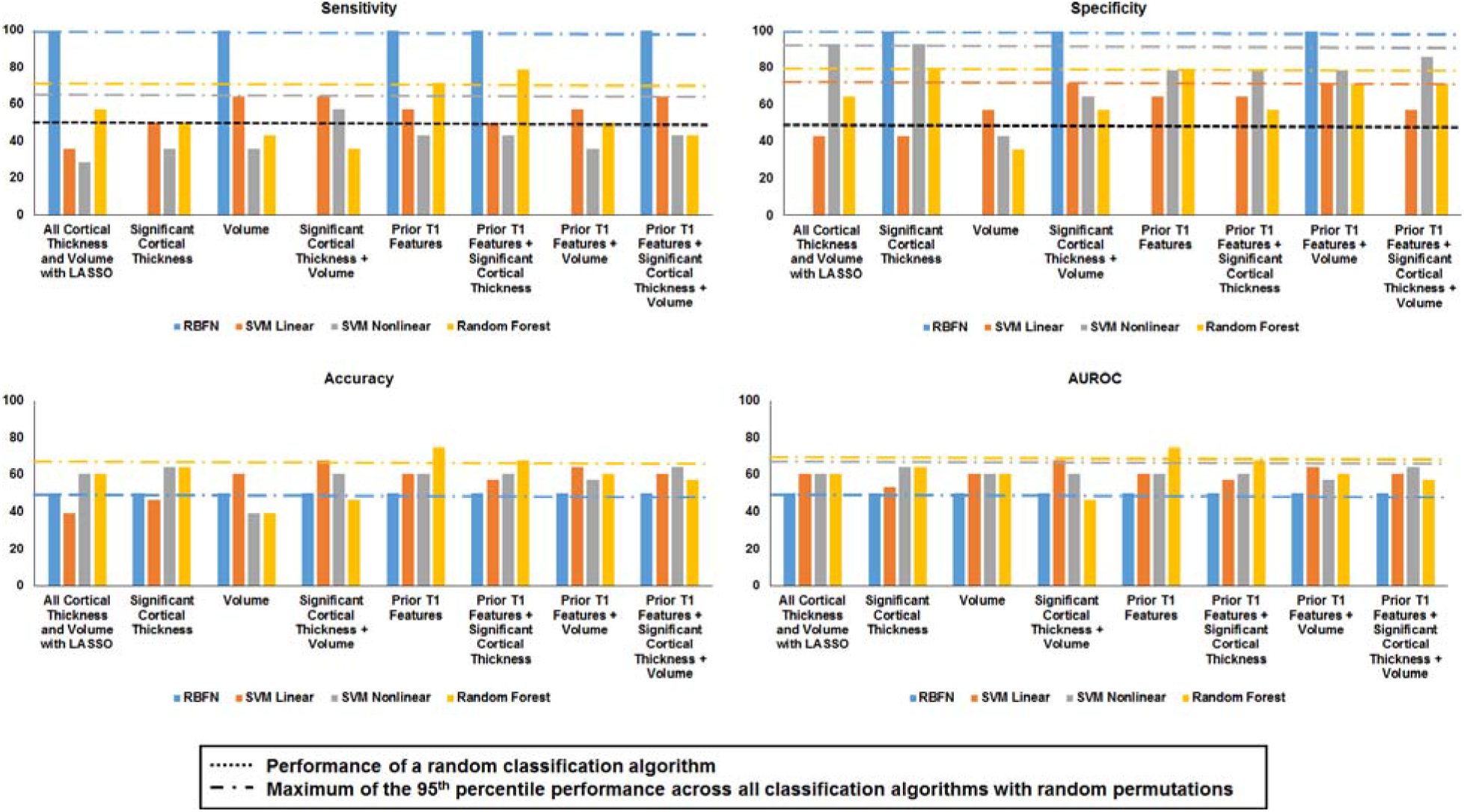
Sensitivity, specificity, accuracy, and area under the receiver operating curve (AUROC) is shown for each machine-learning (ML) algorithm (RBFN: blue, Linear SVM: orange, Nonlinear SVM: gray, and random forest (yellow) for various features. Black dotted lines represent the performance of any random classifier and dotted-dash lines represent the 95^th^ percentile of the benchmark measure for the respective ML algorithm across the various combination of the feature set and are shown in the same colors as the bar-plot.

**Table 3:**
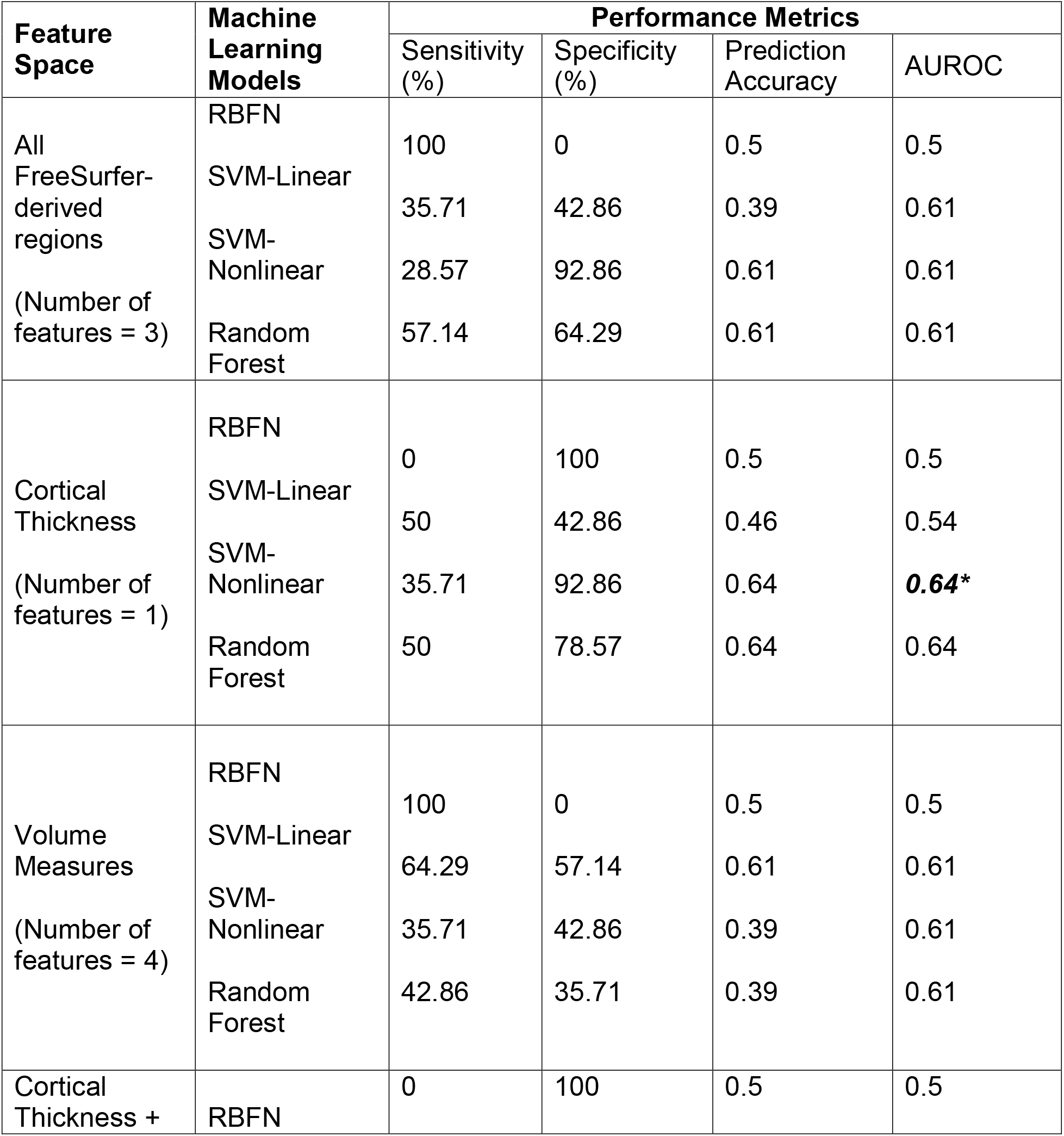
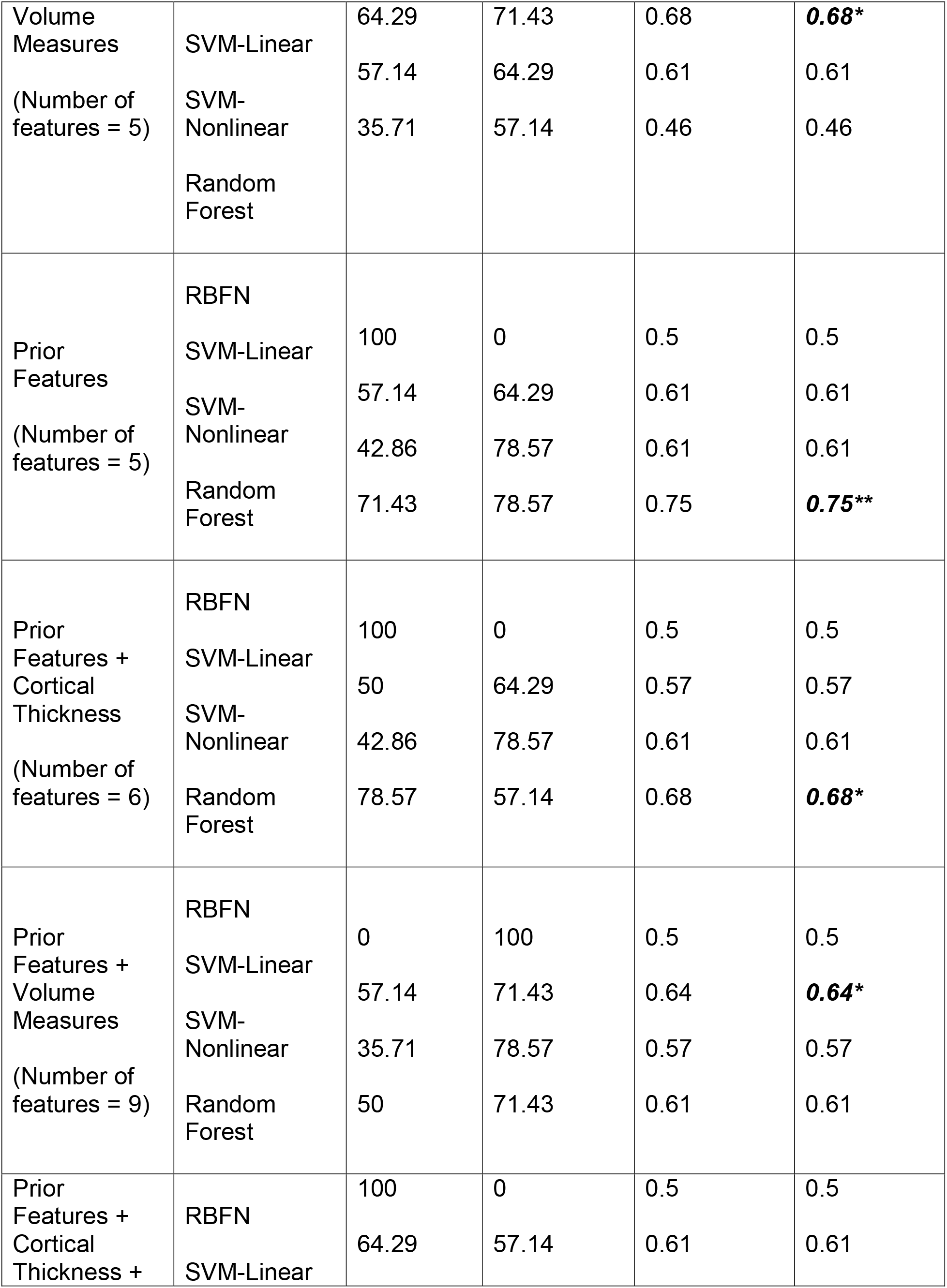
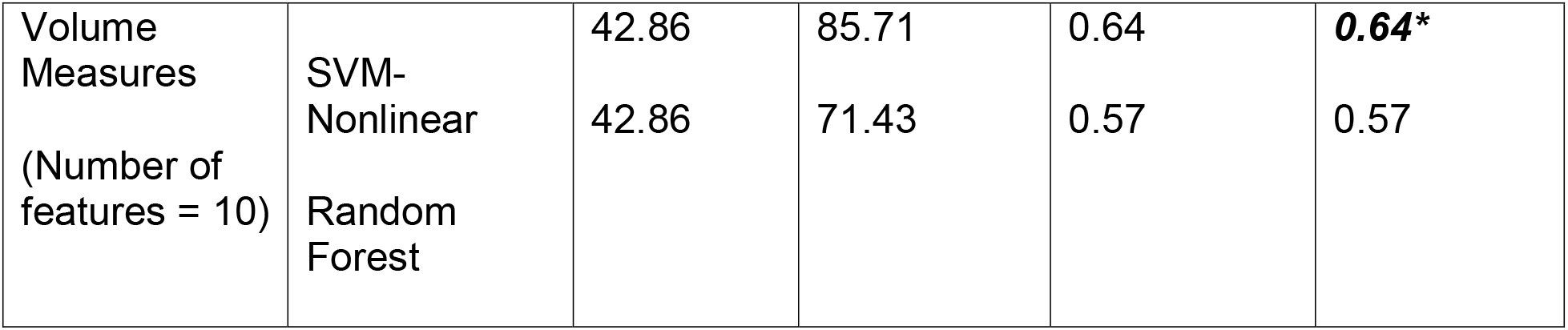
Performance metrics for the various machine-learning models with the associated features are tabulated. AUROC: Area under the receiver operating characteristic; RBFN: Radial Basis Functional Networks; SVM: Support Vector Machine. AUROC results at the 95^th^ percentile are shown in bold with *; AUROC results better than the 95^th^ percentile are shown in bold with **

#### (a) All FreeSurfer-derived cortical thickness and volume

The features identified by LASSO were the temporal-pole volume of the left hemisphere, left hemispheric thalamic volume, and volume of the brain stem. Various ML algorithms were only optimized on these three reduced features. All except RBFN demonstrated an AUROC of 0.61 that was significantly (p_corr_>0.05) less than the benchmark null distribution of 0.68. RBFN had an AUROC of 0.5 which is exactly equal to a random prediction.

#### (b) Regions with only significantly different cortical thickness between the groups

Various ML algorithms were optimized on only one feature, namely cortical thickness of the right cuneus. SVM with the nonlinear kernel (RBF) had a null distribution of AUROC of 0.64 which was at the benchmark significance (p_corr_=0.05). None of the other algorithms achieved a significant AUROC under the null distribution.

#### (c) Only utilizing the volume of the regions showing a correlation between exposure to RHI and cognitive scores

Various ML algorithms were optimized on the following four features: volume of the left fusiform gyrus, GMD cluster predominantly encompassing bilateral thalamus, bilateral putamen, and bilateral caudate, WMD cluster predominantly encompassing the right paracentral lobule, and WMD cluster predominantly encompassing left superior frontal gyrus and left precuneus. Except RBFN, all the ML algorithms achieved an AUROC of 0.61. However, all the models had a significantly (p_corr_>0.05) less AUC as compared to the benchmark null distribution AUC of 0.68. RBFN had an AUROC of 0.5 which is exactly equal to a random prediction.

#### (d) Combination of the regions with significantly different cortical thickness between the groups and volume of the regions showing a correlation between exposure to RHI and cognitive scores

Various ML algorithms were optimized on the following five features: cortical thickness of the right cuneus, volume of the left fusiform gyrus, GMD cluster predominantly encompassing bilateral thalamus, bilateral putamen, and bilateral caudate, WMD cluster predominantly encompassing the right paracentral lobule, and WMD cluster predominantly encompassing left superior frontal gyrus and left precuneus. Only the linear-SVM model achieved an AUROC of 0.68 that was at the benchmark significance (p_corr_=0.05) under the null distribution. None of the other algorithms achieved a significant AUROC under the null distribution.

#### (e) A Priori T1-derived measures

Various ML algorithms were optimized on the following five *a priori* T1-derived features: left cerebellum white matter volume, right hemisphere cortical volume, left thalamus volume, right pallidum volume, and cortical thickness of right rostral-anterior cingulate. Only the random forest algorithm achieved an AUROC of 0.75 which was significantly (p_corr_<0.05) greater than the benchmark null distribution of 0.68. None of the other algorithms achieved a significant AUROC under the null distribution.

#### (f) Combination of a priori T1-derived measures and regions with significantly different cortical thickness between the groups

Various ML algorithms were optimized on the following six features: cortical thickness of the right cuneus, left cerebellum white matter volume, right hemisphere cortical volume, left thalamus volume, right pallidum volume, and cortical thickness of right rostral-anterior cingulate. Only the random forest algorithm achieved an AUROC of 0.68 that was at the benchmark significance (p_corr_=0.05) under the null distribution. None of the other algorithms achieved a significant AUROC under the null distribution.

#### (h) Combination of a priori T1-derived measures and volume of the regions showing a correlation between exposure to RHI and cognitive scores

Various ML algorithms were optimized on the following nine features: volume of the left fusiform gyrus, GMD cluster predominantly encompassing bilateral thalamus, bilateral putamen, and bilateral caudate, WMD cluster predominantly encompassing the right paracentral lobule, WMD cluster predominantly encompassing left superior frontal gyrus and left precuneus, left cerebellum white matter volume, right hemisphere cortical volume, left thalamus volume, right pallidum volume, and cortical thickness of right rostral-anterior cingulate. Only SVM-linear achieved an AUROC of 0.64 that was at the benchmark significance (p_corr_=0.05) under the null distribution. None of the other algorithms achieved a significant AUROC under the null distribution.

#### (i) Combination of a priori T1-derived measures, regions with significantly different cortical thickness between the groups, and volume of the regions showing a correlation between exposure to RHI and cognitive scores

Various ML algorithms were optimized on the following 10 features: cortical thickness of the right cuneus, volume of the left fusiform gyrus, GMD cluster predominantly encompassing bilateral thalamus, bilateral putamen, and bilateral caudate, WMD cluster predominantly encompassing the right paracentral lobule, WMD cluster predominantly encompassing left superior frontal gyrus and left precuneus, left cerebellum white matter volume, right hemisphere cortical volume, left thalamus volume, right pallidum volume, and cortical thickness of right rostral-anterior cingulate. Only SVM-nonlinear achieved an AUROC of 0.64 that was at the benchmark significance (p_corr_=0.05) under the null distribution. None of the other algorithms achieved a significant AUROC under the null distribution.

No graph-theoretical measures, either alone or in combination with other features, were utilized with ML models as there was neither a significant difference observed between any graph-theoretical measures between the groups, nor *a priori* hypothesis can be generated about any graph-theoretical measures that were expected to show a coordinated variation in cortical thickness in our cohort.

## 4. DISCUSSION

This study shows: (i) small but statistical significant differences in cortical thickness in the right cuneus between impaired and nonimpaired boxers but no correlation between cortical thickness and exposure to fighting or cognitive scores in any group; (ii) statistically robust correlation between GMD encompassing caudate, putamen, and thalamus and exposure to fighting but no GMD regions with robust significant group effects; (iii) statistically robust correlation between WMD cluster encompassing right paracentral lobule and psychomotor speed in impaired boxers, and statistically robust correlation between WMD cluster encompassing left superior frontal gyrus and left precuneus lobule and psychomotor speed in nonimpaired boxers but no WMD regions with robust significant group effects; (iv) statistically robust correlation between volume of the left fusiform gyrus and number of fights in nonimpaired boxers; (v) neither a statistically robust effect of coordinated variation in cortical thickness due to RHI between the groups nor a robust correlation between any graph-theoretical network properties and exposure to fighting or cognitive scores in any group; and (vi) that *a priori* hypothesis-driven T1-MRI derived cortical thickness and volumetric measures can reliably identify impaired and nonimpaired active professional boxers when compared to measures derived from regression-based analysis.

### 4.1: Cortical thickness and its role in RHI

Reduced cortical thickness has been shown in participants with mild traumatic brain injury (mTBI) such as professional soccer players (Koerte et al., 2016), motor vehicle collision (Wang et al., 2015), veterans with post-traumatic stress disorder (Lindemer et al., 2013) and early trauma (Corbo et al., 2014), and blast-induced mTBI (Tate et al., 2014). However, cortical thickness studies in boxers are relatively scarce. Contrary to a recent study by Hart et al (Hart et al., 2017), this study found lower cortical thickness in the right cuneus in impaired boxers. Our findings are in agreement with a couple of studies with mTBI participants (Govindarajan et al., 2016; Santhanam et al., 2019) where cortical thinning in cuneus has been reported due to mTBI. The volume of the cuneus is affected with both memory recall and working memory due to TBI (Spitz et al., 2013). However, this study did not reveal any correlations with exposure to fighting or cognitive scores. The absence of any significant statistical difference in the correlation of cortical thickness with cognitive scores or exposure to fighting between the groups suggests a complex and heterogeneous pattern of the role of cortical thickness in participants exposed to RHI. Further investigation into cortical thinning in cuneus due to RHI is warranted, preferably using a longitudinal dataset from the same participants.

### 4.2. VBM and its role in RHI

Depending on the severity and the type of injury, RHI has been shown to produce a cascade of both regional and global brain atrophy (Bigler, 2013; Eierud et al., 2014; Gooijers et al., 2013; Inga K Koerte et al., 2016; Koerte et al., 2015; Langlois et al., 2006). However, similar to our previous findings (Mishra et al., 2018), in our cohort of active professional fighters, we did not find any global or regional VBM differences between our current cohort of impaired and nonimpaired boxers. We speculate that regional or global atrophy due to RHI may lack both sensitivity and specificity for a significant group effect earlier in the pathological process, possibly due to the heterogeneous impact of brain injuries and the inherent repair mechanism of individual boxers.

However, consistent with our previous findings, we did see a cluster encompassing GMD of the bilateral thalamus correlating with years of professional fighting (Bernick and Banks, 2013; Mishra et al., 2018). These findings are in agreement with our previous findings that the thalamus is the most sensitive brain region affected due to RHI (Bernick et al., 2015). This study also showed a robust distinct correlation between psychomotor speed and WMD of paracentral lobule, precuneus, and superior frontal gyrus in our cohort which is in agreement with selective deficits in WMD due to traumatic axonal injury (Warner et al., 2010).

Overall, VBM analysis suggests that though regional or global brain atrophy is not evident in active impaired and nonimpaired boxers, there is a distinct pattern of correlation between both GMD and WMD and exposure to fighting and neuropsychological scores. Future longitudinal investigation utilizing VBM analysis may reveal if there is preferential atrophy of cortical or subcortical regions that are not only different between impaired and nonimpaired active boxers but also correlated with exposure to fighting and cognitive scores.

### 4.3: Graph-theoretical analysis and its role in RHI

This study neither found any statistically significant group-level differences between our cohort of active impaired and nonimpaired boxers nor any correlation with exposure to fighting or cognitive scores which is contradictory to previous graph-theoretical analysis studies in participants with neurodegenerative brain disorders (He et al., 2008; Wang et al., 2016) and TBI (Proessl et al., 2020). The absence of any significant group-level statistical difference combined with the absence of correlation of any graph-theoretically derived measures with cognitive scores or exposure to fighting between the groups further bolsters our initial speculation that RHI induces a complex and heterogeneous modification in cortical thickness in participants exposed to RHI. However, a longitudinal investigation into coordinated changes in cortical thickness due to RHI is warranted, preferably using a longitudinal dataset from the same participants.

### 4.4: Individual prediction of impaired and nonimpaired active professional boxers

Despite forcing double-dipping on our feature selection from regression-based methods, our exploratory analysis suggested that none of the ML algorithms was able to confidently identify impaired active professional boxers. At the very best, any individual classifier was at the 95^th^ percentile, which was the threshold used to select the features. Furthermore, a clear preference of overfitting the prediction accuracy was revealed, especially when the RBFN model with the entire FreeSurfer derived measures was utilized, despite a well-curated and balanced sample set. Our findings are against the belief that double-dipping leads to an inflated prediction accuracy (Bishop, 2006; Demirci et al., 2008). Rather, our findings suggest that ML models must utilize an independent testing dataset and test the benchmark accuracy against the random group assignments as suggested by Arbabshirani et al. (Arbabshirani et al., 2017).

Although only a subset of previously identified hypothesis-driven feature sets (Mishra et al., 2017) was utilized in this study, we observed a greater-than-chance (p_corr_<0.05) prediction accuracy of 75% to identify impaired active professional boxers. Utilizing such hypothesis-driven features to drive the classifier bolsters our previous finding and suggests that though the hypothesized brain regions might not show statistical regional or voxel-based differences between the groups, the nonlinear combination of *a priori* brain regions might be better suited for individual prediction in neuroimaging studies.

### 4.5: Limitations and future work

This study must be discussed in light of the following limitations. First, any topological difference in brain network architecture due to professional fighting is currently unknown since the data from controls were not acquired for this study. Second, similar to Gao et al (Gao et al., 2014), a multimodal MRI study in cognitively impaired active boxers might shed light on the combined anatomical and functional connectivity modifications in the brain due to RHI. Third, future studies could combine the T1-MRI-derived measures identified in this study with the dMRI-derived measures in the same participants, as obtained in the previous study (Mishra et al., 2017), to investigate the increase in the prediction accuracy of identifying impaired and nonimpaired active professional boxers. Fourth, the recent improvement in the deep-learning algorithm (Thomas et al., 2019) may be utilized to further observe the improvement in prediction accuracy as compared to conventional ML algorithms used in this study. Fifth, the ML results are only exploratory in nature. Though the testing dataset is independent of the training dataset, the sample size in the testing dataset is small (n-14 in each group), and hence the results of this study need to be confirmed with a bigger dataset. Sixth, correlation between morphometric changes and role of concussions or other medical trauma associated with exposure to fighting, presence of psychiatric disorders, drug and pain medication use, and drug-use disorder should be evaluated in future studies as these measures are also associated with cognitive impairments due to RHI. Lastly, future longitudinal studies with larger samples and more comprehensive characterization of impaired and nonimpaired boxers will allow for more robust investigation of the predictive power of the brain regions identified in this study to monitor cognitively impaired active professional boxers as this study may just be looking at cognitive differences and the boxers may not be on a neurodegenerative trajectory.

## 5. CONCLUSION

In conclusion, this study revealed that T1-weighted MRI measures can reveal significant morphometric differences in cortical thickness and volumetric measures that are correlated with low cognitive scores and exposure to fighting between impaired and nonimpaired active professional boxers. Exploratory analysis with various ML algorithms further revealed that despite the presence of such group-level morphometric differences, utilizing such regression-driven group-level analysis might not identify impaired and nonimpaired active professional boxers. However, *a priori* hypothesis-driven T1-derived cortical thickness and volumes of certain brain regions are better capable to identify impaired and nonimpaired active professional boxers.

## ACKNOWLEDGEMENTS

We would like to extend our sincere thanks to all the participants of the study, various research coordinators and MRI technologists without which the study would not have been completed. We would also like to thank Dr. Mark Lowe and Dr. Wanyong Shin from Cleveland Clinic for their assistance in setting up the MRI protocols at our center. The funding sources have no role in the study design, data collection, data analysis, interpretation, or writing of this article. All the authors had unrestricted access to the data in the study and had a final decision to submit it for publication.

